# Tripleknock: predicting lethal effect of three-gene knockout in bacteria by deep learning

**DOI:** 10.1101/2025.07.31.667916

**Authors:** Peter X. Geng, Jiaheng Hou, Jinyuan Guo, Xiaoqing Jiang, Huaiqiu Zhu

## Abstract

Investigating the lethal effect of multi-gene knockout is essential for discovering novel antibiotics targets and metabolic engineering. Unlike single genes or gene pairs, three-gene combinations involve more intricate interactions, making experimental screening time-consuming. Computational methods, particularly Genome-scale metabolic Model (GEM)-based Flux Balance Analysis (FBA), requires constructing new GEMs from experimental data, limiting its use for new species. Moreover, using FBA for three-gene knockout screening could take several years. Therefore, a faster and GEMs-independent approach is needed to facilitate genome-wide three-gene knockout screening. Here, we introduce Tripleknock, for predicting the lethal effects of three-gene knockouts. Tripleknock was trained using whole-genome data from *Escherichia coli* K-12 MG1655, and three-gene knockout simulations using FBA. The model uses a threshold of 90% reduction in cell growth to define lethal effect as the prediction output. Compared to FBA, Tripleknock achieves predictions approximately 20 times faster, reaching an average cross-species F1 score of 0.77 on six pathogenic species within the *Enterobacteriaceae* family. For closely related species such as pathogenic *E. coli* and *Shigella*, Tripleknock reaches F1 scores exceeding 0.83. To our knowledge, Tripleknock is the first end-to-end model for predicting lethal effects of three-gene knockout in bacteria.

**Data availability:** Tripleknock is publicly available at: https://github.com/Peneapple/Tripleknock

## Introduction

Gene knockout is a widely-used method to investigate the relationship between genotype and phenotype. In metabolic engineering, gene knockouts can generate bacterial strains with increased production of specific metabolites^1^, while in antibiotic design, gene-knockout screening can facilitate the discovery of novel drug targets^2^. Single-gene knockout studies have paved the way for identifying genes important for survival and growth. These genes are critical for maintaining basic cellular functions, such as metabolism, DNA replication, and cellular integrity. Absense of essential genes will disrupt cellular processes, typically halting growth or leading to cell death^3–5^. The identification of essential genes has been primarily achieved through experimental methods such as gene editing, RNA interference, and CRISPR/Cas9^6,7^. In addition to experimental methods, computational approaches have also been developed to predict essential genes across species based on genomic data^8,9^.

Compared with single-gene kockout, two-gene knockout can reveal novel genetic interactions that are not evident in single-gene knockout studies. Among these interactions, Synthetic Lethal (SL)^10^ and Synthetic Rescue (SR)^11^ are two important types of gene interaction. SL is defined as when the knockout of both genes results in a severe loss of cell growth or even cell death, while the knockout of either gene does not lead to this inactivation, indicating that both genes are functionally interdependent. On the other hand, SR refers to a situation where the inhibition of one gene (either gene *A* or gene *B*) alone reduces cell growth, but the simultaneous inhibition of both genes rescues the phenotype, thus allowing the cell to survive. SR is also a potential new mechanism in cancer drug resistance^12–14^. These interactions provide valuable insights into gene dependencies and redundancies in cellular networks. Algorithms and experimental methods for understanding SL and SR effects are also reported^15–17^.

Expanding to three-gene knockouts, the complexity of genetic interactions increases even further compared to two-gene interactions. Exploring three-gene knockout combination is important and has shown promise in metabolic engineering and therapeutic interventions. For example, a three-gene knockout has also been reported to enhance polyhydroxyalkanoate (PHA) production in *E*. coli^18^. These interactions could result in unexpected phenotypic effects, and understanding them may provide new avenues for therapeutic interventions. For pathogenic species, investigating the lethal effect of multi-gene knockout could facilitate the finding of new antibiotics targets. However, multi-gene knockout remains a significant challenge, for the difficulties of accessing sufficient experimental data. And the lack of experimental data limits the development of computational methods. Fortunately, GEMs coupled with FBA^19,20^ provide reliable results for simulating cell growth when genes are knocked out. But, although it’s the most popular simulation methods in the past over 20 years, it faces limitations when applied to multi-gene knockout. First, the construction of GEMs for each species is labor-intensive and time-consuming^21–24^. Even if the new species is closely related to a species that already has GEM, a new GEM must be built de novo, which makes it difficult to generalize the results of gene knockout across related species. Second, GEMs only include approximately one-third of genes in the genome, which means the searching space is largely limited. Recent advancements in deep learning have shown promise in predicting gene knockout effects without relying on GEM construction^25–27^. Given these constraints, an accurate, GEM-independent and user-friendly prediction model for multi-gene’s lethal effects would be a powerful tool for biologists who are working on metabolic engineering or multi-target antibiotics finding.

In response to these challenges and inspired by deep learning models, we developed Tripleknock, to predict three-gene knockout effects in *E. coli* related species based on whole-genome sequence. Cross-species prediction demonstrated the power of Tripleknock in predicting lethal three-gene knockouts on six pathogenic species. This approach offers a solution to overcome the current limitations in studying the lethal effects of multi-gene knockout, making it an assistive tool for future research in antibiotics finding.

## Results

### Overview of Tripleknock’s Algorithmic Architecture

The Tripleknock workflow consists of three main components (Fig. 1). First, a two-stage pre-trained autoencoder is used for genome compression (Fig. 1a). This tandem autoencoder model is designed to process genomes with triple gene knockouts and sequentially reduces both the dimensionality of gene features and the number of genes, resulting in an approximately 1,000-fold reduction in genome matrix size. Second, phenotype labels are generated using FBA, where all feasible three-gene knockout combinations in the GEM of *E. coli* K-12 MG1655 are simulated and categorized based on their predicted growth outcomes (Fig.1 b). Third, a classification model based on a multilayer perceptron (MLP) is used to predict the viability of triple knockout genotypes (Fig. 1c). This classifier receives a combined feature consisting of the compressed genome and the triple knockout genes, and processes it through fully connected layers to produce a binary prediction. Both the autoencoder and the MLP models are available for direct download and application for users.

**Fig. 1.**
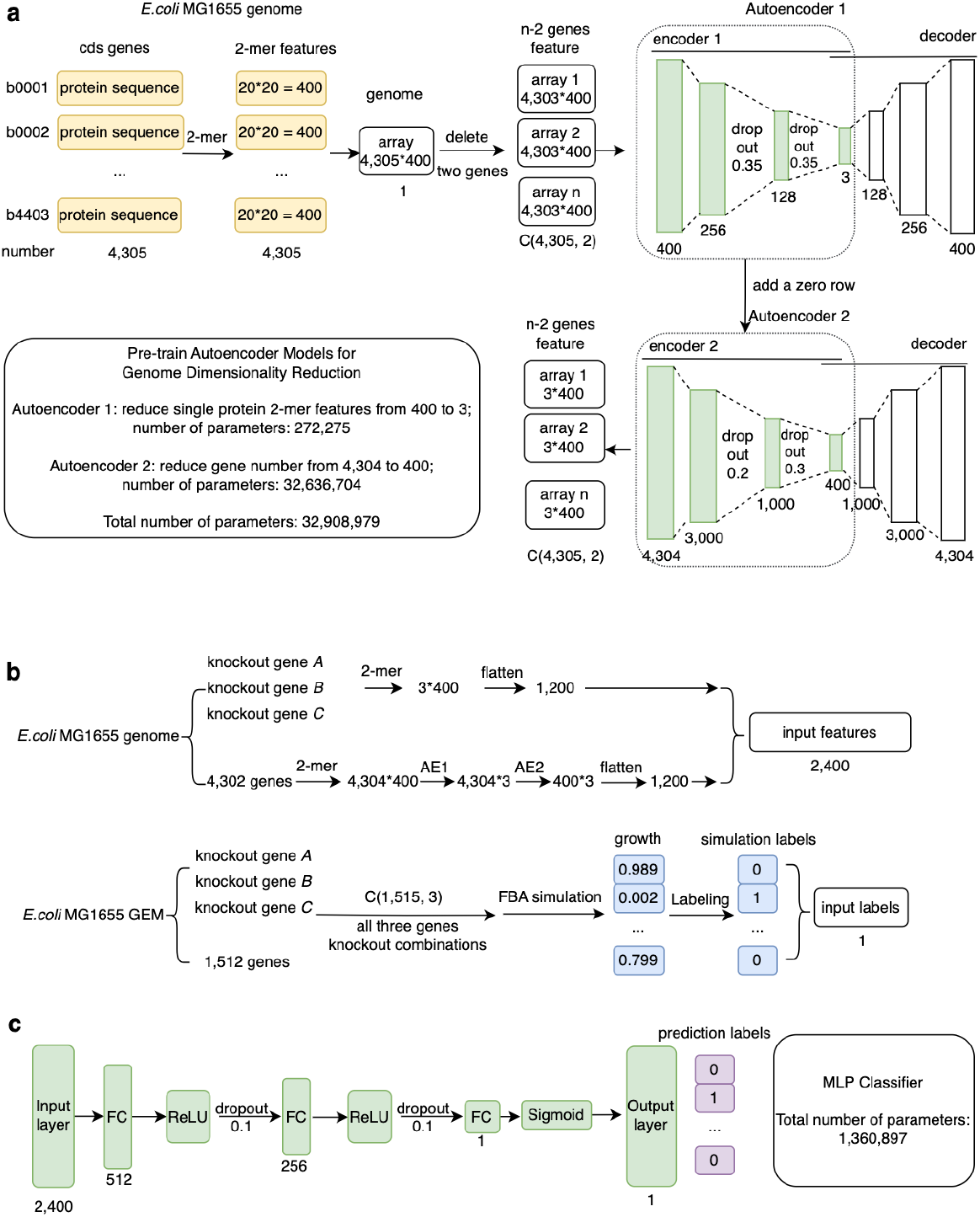
Workflow of Tripleknock. **a**, Construction of 2-mer encoded protein feature matrices and genome-level compression via a two-stage autoencoder pipeline. **b**, Feature engineering and three-gene knockout simulation by FBA. **c**, Predictions using concatenated features of knocked-out genes and genomes, done by MLP-based binary classification of cell growth. FC, fully connected layer. Yellow boxes refers to feature engineering, green boxes indicates the layers in three sub architectures, blue boxes indicates the labeling method, and the purple boxes is the final binary output of Tripleknock.

We plotted the loss function of the three sub-models during training (Supplementary Fig. 2). Autoencoder 1 (AE1) demonstrates rapid convergence. A horizontal dashed green line indicates the theoretical background noise level (1/400 = 0.0025), which reflects the average frequency of a 2-mer assuming uniform distribution. The model’s reconstruction error is significantly below this threshold, indicating that the compressed embedding preserves meaningful biological signal beyond random noise (Supplementary Fig. 2a). Autoencoder 2 (AE2) achieves extremely low reconstruction error. This reflects the high regularity and strong redundancy in the genome-wide compressed feature space (Supplementary Fig. 2b).

For MLP training, we estimated that the time required for generating and storing all files during training will be approximately 70 days; considering a 10-fold cross-validation, this will increase to around 700 days. Therefore, only random 5% and 10% subsets were used to train the model (Supplementary Fig. 2c, 2d). In both cases, the loss steadily decreased and stabilized. We performed 10-fold cross-validation on both 5% and 10% subsets. In both cases, the model achieved an exceptionally high average Area Under the Curve (AUC) score of 0.9998. Notably, the training on 5% and 10% subsets yielded similar training curves, justified the use of subsampling method.

The time of data preparation (FBA simulation) and trainig of sub-models within Tripleknock framework is collected (Table 1).

**Table 1.** Data collection and training time of models in Tripleknock Step Time AUC.

| Step | Time | AUC |
| --- | --- | --- |
| **FBA simulation** | ~100 days (100-threads) | \ |
| Sub-model training |  |  |
| Autoencoder 1 | 8.15 | \ |
| Autoencoder 2 | 19.72 | \ |
| MLP 5% | ~5 | 0.9998 (10-fold) |
| MLP 10% | ~10 | 0.9998 (10-fold) |
| **Total training time** | ~30-40 | \ |

### Cross-species prediction

We evaluated the cross-species generalizability of the Tripleknock framework by applying it to six pathogenic bacterial species within the *Enterobacteriaceae* family. For each species, 20,000 unique three-gene knockout combinations were simulated using FBA, with each simulation repeated three times to generate a total of 60,000 combinations per species. The 10% training version of Tripleknock was then used to predict lethal effect based on the gene ID and annotated genome sequences. The resulting average F1 scores for each species were mapped onto a phylogenetic tree (Fig. 2a), demonstrating that Tripleknock achieved an overall cross-species average F1 score of 0.77, with particularly strong performance observed in *E. coli* and *Shigella*, where F1 scores exceeded 0.83. To investigate the impact of training data size on predictive performance, we compared classification metrics (Accuracy, Precision, Recall, and F1-score) across models trained with 5% and 10% of the data (Fig. 2b). The differences were not statistically significant according to the Wilcoxon signed-rank test, suggesting that 10% training data is sufficient for robust cross-species generalization. Accordingly, all subsequent interpretability analyses were conducted using models trained on 10% of the data, and the version of Tripleknock released for public use is likewise based on this setting. In terms of computational efficiency, Tripleknock delivers predictions consistent with FBA outputs approximately 20 to 30 times faster when executed on an L4 GPU within the Google Colab environment. Testing on Colab also suggested the usage and capability of Tripleknock for users.

**Fig. 2.**
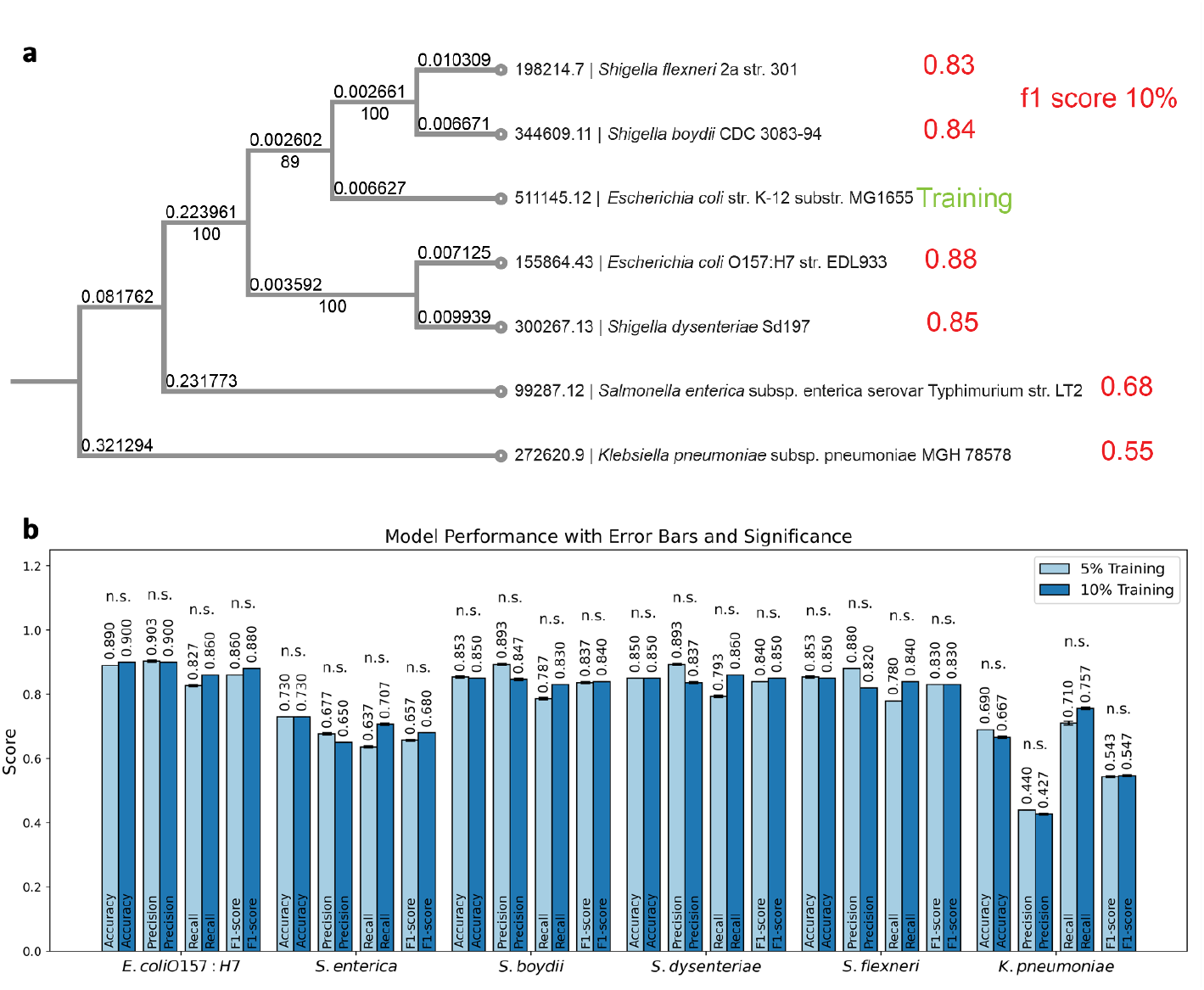
Cross-species prediction. **a**, Phylogenetic tree of the seven selected species within the *Enterobacteriaceae* family. Branch lengths are indicated above the grey lines, and confidence values are shown below. Genome IDs are displayed to the right of each node. Average F1 scores for each species are annotated in red. The training data usage is 10%. **b**, Performance of Tripleknock trained on 5% and 10% of *E. coli* K-12 data and evaluated on six pathogenic species. Error bars represent the standard error of the mean. The average values are labeled above each bar. Statistical significance between the two training ratios for each species was assessed using the Wilcoxon signed-rank test.

### Model interpretation

Model interpretability was evaluated on three representative test species based on their prediction performance: the species with the highest F1 score, the one with the lowest F1 score, and a randomly selected *Shigella* species. In the hexbin volcano plot (Fig. 3a, 3b, 3c). The well-performing species (*E. coli* O157:H7, iZ_1308) exhibits clearer separation and stronger significance, indicating more distinguishable informative features. Density plots of top features in positive and negative samples (Fig. 3d, 3e, 3f) show distributions of the top three features. Feature enrichment plots (Fig. 3g, 3h, 3i) assess how top-ranked features are enriched in the first 1,200 dimensions, which correspond to features derived from the triple-knockout genes themselves. In *E. coli* O157:H7, enrichment is concentrated in early ranks, suggesting model decisions are heavily influenced by the knocked-out gene features. In contrast, *Klebsiella pneumoniae* (iYL1228) drops steeply at the early stage, indicating weaker signal recognition. Tripleknock’s interpretability correlates with predictive accuracy—species with better performance exhibit clearer feature-level separation and stronger signal enrichment. This validates the model’s decision logic and confirms its effectiveness on phylogenetically or functionally similar species.

**Fig. 3.**
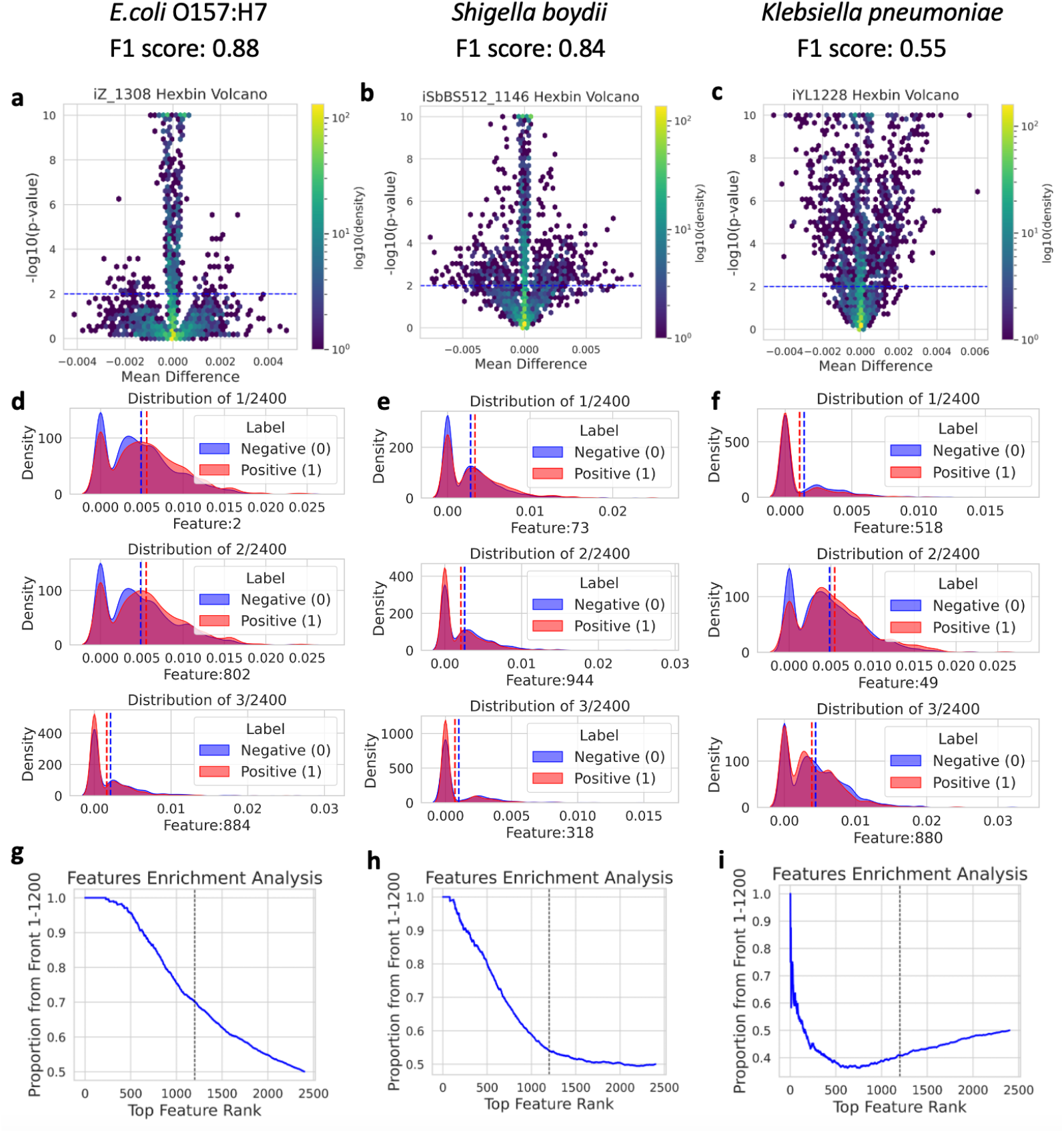
Feature analysis in three test species. **a-c**, Hexbin density volcano plot. The colorbar indicates the number of samples. The x-axis represents the mean difference between the two groups, the y-axis shows the −log10-transformed p-value from a Welch’s t-test. The blue dashlines indicate the p-value 0.01. **d-f**, The density distribution estimation of the top 3 features in positive samples and negative samples in test data, with vertical dashed lines indicating the mean value of each class. **g-i**, Feature enrichment analysis in the test data, with a black dashed line indicating the mid-point of 1,200.

## Discussion

The lethal effect of multi-gene knockout screening is very useful for finding antiobiotics target and metabolic engineering. Two significant challenges in the utilization of GEM-based FBA are the time-consuming screening of multi-gene knockouts and GEM-dependency. In light of this inefficiency, and taking lethal effect of gene knockout as an example, we conducted a comprehensive screening of all possible three-gene knockouts in *E*.*coli* K-12 and collected the cell growth data after gene knockout. We developed a deep learning algorithm named Tripleknock that successfullly predicted the cell growth after three-gene knockouts across six pathogenic species. The model interpretability analysis assessed how features contribute to predictions. Feature enrichment analysis across species revealed substantial variability, indicating that the interaction between the three knockout genes and the genome plays a critical role in model generalization. As evolutionary distance increases, the functional and feature-level differences among the same three knockout genes also increase, which in turn hampers the model’s cross-species predictive capacity. These findings suggest a future direction that a deeper understanding of gene–genome interaction is essential to improve the model’s transferability across species^28^.

During the feature extraction phase, we tried to integrate DNA sequence and GEM topological information. However, feature ablation analysis revealed that only protein sequence data yields the best results. We manage to make predictions without depending on GEM. The use of GEM presents a challenge, as constructing a high-quality GEM model for a new species is not straightforward. We build Tripleknock to function end-to-end and is friendly for biologists who are not familiar with coding, solely based on an annotated genome file, without relying on external or supplementary materials. Some other questions will also be considered. For instance, this study employs a binary classification approach using solely sequence data, which may be inadequate for more complex predictive tasks. Moreover, our prediction does not incorporate experimental metabolic data, such as multi-omics datasets. It is estimated that integrating these diverse data sources could enhance the accuracy of cross-species predictions. Sequence similarity represents just one of many conserved features across different species; other factors, such as metabolic network-based information^29^, also contribute significantly to cross-species prediction. Therefore, both sequence conservation and metabolic structural conservation are critical for comprehensive cross-species analysis.

In summary, we developed the first end-to-end deep learning-based three-gene knockout model Tripleknock, which successfully predicts the lethal effect of three-gene knockout in the *Enterobacteriaceae* family. Our model is an effective and user-friendly tool for biologists who are working on multi-target antibiotics finding or metabolic engineering.

## Methods

### Computational environment

The FBA simulation and model training were performed on a server running an Ubuntu 18.04.2 LTS system. The hardware included 4 Intel(R) Xeon(R) Gold 6252 CPUs operating at 2.10 GHz, and 2 NVIDIA A100-PCIE-40GB GPUs. The software environment was based on Python 3.8.19, CUDA Version 11.7. Cross-species evaluations were conducted on Google Colab using an L-4 GPU. Model interpretability was conducted on Google Colab using a CPU. No software version-dependent errors were encountered, demonstrating the robustness and portability of Tripleknock.

### Datasets

*E.coli* K-12 MG1655, a non-pathogenic strain, was used for training. For testing, we selected one *E. coli* strain, one *Salmonella enterica*, three *Shigella* species, and one *Klebsiella*, all of which are pathogenic. These seven species belong to the *Enterobacteriaceae* family (Table 2). The GEMs were downloaded from the Biochemical, Genetic and Genomic knowledgebase (BiGG), a database of high-quality GEMs^30^. The protein Coding DNA Sequences (CDS) for each species were downloaded from the National Center for Biotechnology Information (NCBI) in FAST-All (FASTA) Protein format. A horizontal bar plot shows the number of genes annotated in the genome (CDS genes) versus those included in the GEMs across seven species (Supplementary Fig. 1). All of the distributions of protein length of the seven species are right-skewed, indicating that most proteins are relatively short, typically under 600 amino acids. A long tail exists, representing a small number of much longer proteins, some exceeding 2,000 amino acids.

**Table 2.** Selected bacteria in training and test set.

| Organism | BiGG | NCBI | BV-BRC | Pathogenic |
| --- | --- | --- | --- | --- |
| *E. coli* str. K-12 substr. MG1655 | iML1515 | NC_000913.3 | 511145.12 | No |
| *E. coli* O157:H7 str. EDL933 | iZ_1308 | AE005174.2 | 155864.43 | Yes |
| *Salmonella enterica* subsp. enterica<br>serovar Typhimurium str. LT2 | STM_v1_0 | AE006468.2 | 99287.12 | Yes |
| *Shigella boydii* CDC 3083-94 | iSbBS512_1146 | CP001063.1 | 344609.11 | Yes |
| *Shigella dysenteriae* Sd197 | iSDY_1059 | CP000034.1 | 300267.13 | Yes |
| *Shigella flexneri* 2a str. 301 | iSF_1195 | AE005674.2 | 198214.7 | Yes |
| *K. pneumoniae* subsp. pneumoniae<br>MGH 78578 | iYL1228 | CP000647.1 | 272620.9 | Yes |

The ‘Bacterial Genome Tree’ tool in the Bacterial and Viral Bioinformatics Resource Center (BV-BRC)^31^ was used for building the reference phylogenetic tree for cross-species prediction (Supplementary Table 1). This was used for cross-species prediction, during which we used Tripleknock to receive an annotated genome sequence to make direct three-gene knockout prediction on these related species.

### Pretrained tandem autoencoders

The first process generated all two-gene pairs from a total of 4,305 genes (*G*^*CDS*^), in the *E.coli* K-12 MG1655 genome. The total number of combinations is 9,264,360. The removing of each two-gene combination from *G*^*CDS*^ will lead to an n-2 gene set, which is represented as a matrix whose shape is 4,303 × 400. The AE1 was trained to compress gene feature vectors from 400 dimensions down to 3, and the AE2 reduces the dimensionality of gene number to 400. The training process used all data over 6 epochs, with a batch size of 2,000. The equation (1) is the loss function used for training is Mean Squared Error (MSE) to minimize the reconstruction error between the input and the output.

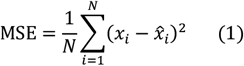

Where:

*N* is the number of input gene features in AE1, the number of genes in AE2.

*x*_*i*_ is the original input.

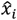 is the reconstructed output from the autoencoder (the predicted feature vector).

To accommodate Tripleknock for potential single-gene knockout on *E.coli* MG1655, we explicitly add one zero vector row to the input matrix (making it 4,304 × 400). This ensures that, when simulating single-gene knockouts for a genome of 4,305 genes, every gene in *E. coli* K-12 MG1655 is represented and the matrix multiplication aligns with the model’s input dimensions. Without adding zero rows, predicting a single-gene knockout would require randomly omitting a gene to satisfy the multiplication rules.

Given a two genes knockout genome matrix 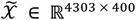, we create an extended matrix 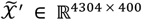 by appending a zero row as equation (2):

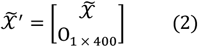

Before training, each feature vector is normalized using min-max normalization to ensure that the values across different genes are on a comparable scale.

For each feature vector 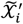, we apply the min-max normalization as equation (3):

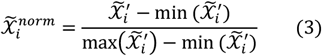

If 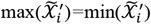, then:

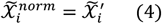

The genome matrix 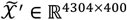 is compressed to 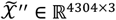 by the encoder of AE1, which then is compressed to 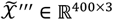 by the encoder of AE2.

Dropout regularization was applied to both autoencoders (0.35 in AE1; 0.2 and 0.3 in AE2). This autoencoder pipeline had a total of ~32.9 million parameters.

### Gene knockout simulation by FBA

We first generated all three-gene combinations form the single gene set of the GEM of *E.coli* K-12 MG1655 (BiGG: iML1515), which has 1,515 genes (manually removed s0001). For each three gene combination, the knockout simulation was simulated using CobraPy^32^ with the default target function by maximizing its biomass. We calculated whether the cell growth is less than 10% of the initial growth by equation (5):

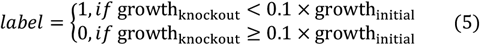

After performing the knockout for all combinations, we proceed with data cleaning by removing infeasible knockout combinations and excluding those that involving the gene *b2092* and gene *b4104*, which are pseudogenes based on the NCBI record (accessed by Mar 09, 2022). The theoretical number of possible three-gene combinations is:

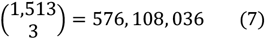

After data cleaning and merging, we obtain 576,107,619 valid combinations, with 417 missing due to infeasible FBA results. Among all valid combinations, 380,596,069 were labeled as 0, and 195,511,550 were labeled as 1. The ratio of positive to negative labels is 1:1.95. The merged data (12.1 GB) was randomly shuffled and then split into 10 parts for training and the next 10-fold cross-validation.

The complete simulation of all three-gene knockout combinations was conducted over a period of two months, utilizing 100 parallel computational threads.

### MLP classifier

For each three-gene combination 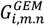, two parts of the features are constructed and concatenated as input for the classifier.

The first part is three-gene 2-mer frequency features. Each gene *g* = {*i, m, n*} has a 2-mer frequency vector of size ℝ^400^, which are concatenated into a single feature vector of size ℝ^1,200^:

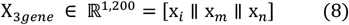

Then, z-score normalization is applied:

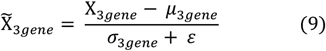

Where *μ* and *σ* denote the mean and standard deviation of the vector, and *ε* = 10^−8^ prevents division by zero. The second part is remaining genome features via dual autoencoders. The remaining genome is defined as:

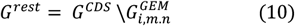

Each gene in *G*^*rest*^ is represented by a 2-mer frequency vector in ℝ^400^, forming a matrix M ∈ ℝ^4,302×400^. To maintain the consistent shape, two zero vectors are added:

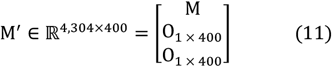

This matrix is passed through a tandem autoencoder pipeline, reducing the dimensionality to:

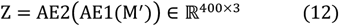

The compressed matrix Z is flattened and normalized:

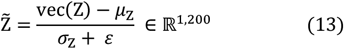

The final input to the MLP is:

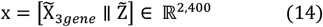

The input x ∈ ℝ^2,400^ is passed to an MLP classifier with sigmoid output, predicting the probability of a lethal 3-gene knockout. The model is trained with Binary Cross Entropy Loss (BCELoss), defined as:

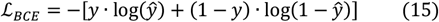

Where:

y is the true label

*ŷ* = MLP(x) ∈ (0,1) is the predicted probability

The model was then trained using the Adam optimizer with a learning rate of 0.001, and trained for 4 epochs, 10% of data was used for testing, 10-fold cross-validation was also performed.

### Model interpretability

For volcano visualization, we calculated the difference in mean values for each feature between positive and negative classes, followed by Welch’s t-test to evaluate statistical significance. The results were visualized using hexbin density plot. A test dataset has 20,000 three-gene combinations, and each gene combination corresponds to 2,400 features. Here we let X_*pos*_ ∈ ℝ^*n*1×2,400^ and X_*neg*_ ∈ ℝ^*n*2×2,400^ be the feature matrices for the positive and negative groups.

We calculate the mean difference:

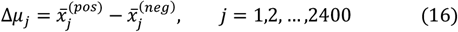

Where 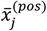 is the mean of feature *j* in the positive group.

Then we do statistical significance by Welch’s t-test:

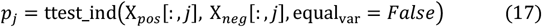

Log-transformed p-values:

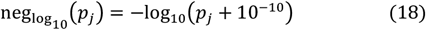

For the density distribution of top-ranked features. After computing the mean difference and statistical significance for each feature, we ranked all features based on their −*log*_10_(*p*) score. The top 3 features were selected for visualization. For each feature, we plotted the probability density distributions across the two classes (label = 0 and label = 1). The distributions were estimated using kernel density estimation (KDE). This visualization highlights the distinct distributional shifts of the most discriminative features.

Here, the mean value for each class is calculated by:

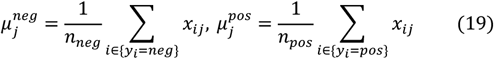

Kernel density estimate:

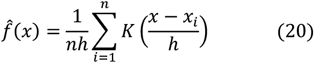

Where n is the sample number, *K* is the Gaussian kernel, and *h* is the bandwidth.

For feature enrichment analysis to investigate whether the top-ranked features were disproportionately concentrated in the first half of the input vector, which corresponds to the region associated with the three target genes. After ranking all features by statistical significance, we calculated, for each rank *i* ∈ {1, …, 2400}, the proportion of the top *i* features that originated from the first 1,200 indices. This proportion was then plotted against the feature rank to reveal potential enrichment trends in the early part of the ranked list.

Here we let *T* = [*t*_1_, *t*_2_, …, *t*_*N*_] be the list of feature indices sorted by significance, where *N=*2,400 is the total number of features. Let *A* = {1,2, …, 1200} denote the index range corresponding to features derived from the three genes.

For each rank *i* = {1,2, …, *N*}, we define the enrichment ratio *r*(*i*) as:

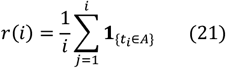

Where:

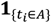 is an indicator function that equals 1 if the *j*-th top-ranked feature belongs to the front half (three gene knockout candidates), and 0 otherwise.

*r*(*i*) represents the cumulative proportion of features from the front part among the top *i* ranked features.

This function is used to quantify how enriched the top-ranked features are within the three-gene feature or the rest of the genome. Under a uniform distribution with no enrichment, the expected value of the ratio *r*(*i*) is approximately 0.5 for all *i*.

## Acknowledgements

This work was supported by the National Key Research and Development Program of China (2021YFC2300300) and the National Natural Science Foundation of China (32070667). Part of the analysis was performed on the High-Performance Computing Platform of Peking University.

## Author contributions

G.P.X. and Z.H. led the research. G.P.X. collected the data, developed the network architecture and training, and conducted the interpretability analysis. H.J., G.J., and J.X. contributed numerous ideas throughout the project.

## Data availability statement

The original code for data collection, which includes three gene knockout FBA simulations, GEM models, and model training using an autoencoder and MLP classifier, is stored on GitHub at the following link: https://github.com/Peneapple/Tripleknock. On the main page of GitHub, we also offer a tutorial for new users unfamiliar with coding to run Tripleknock on Google Colab.

## Supplementary

**Supplementary Fig. 1.**
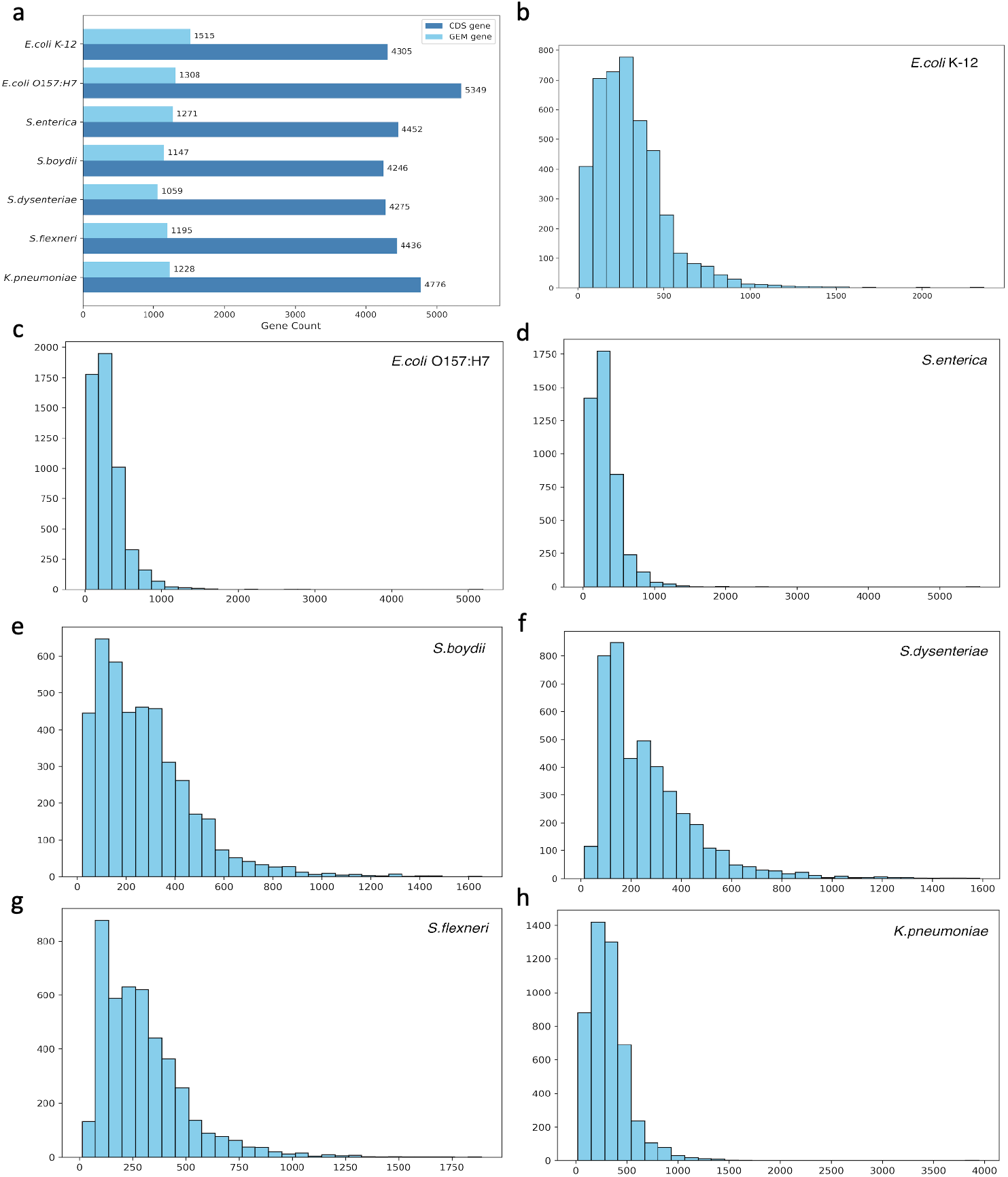
The statistical analysis of the selected seven species. **a**, The number of genes in each genome and GEM. The light blue bars represent the total number of genes in the GEM; the dark blue bars show the number of CDS genes in the genome. **b-h**, The distribution of protein lengths (in amino acids) for seven species. Each panel shows the number (y-axis) of proteins within specified length intervals (x-axis).

**Supplementary Fig. 2.**
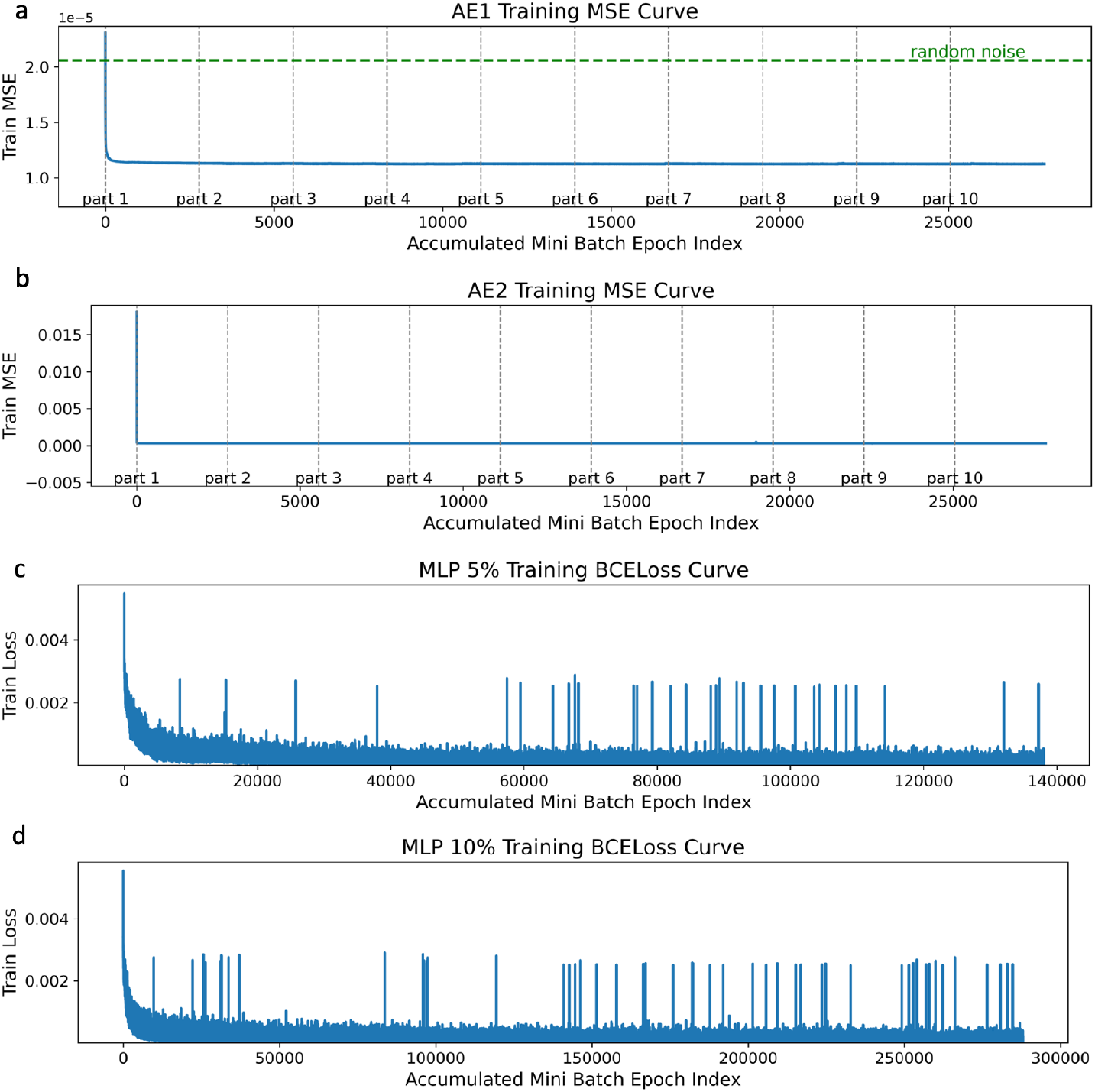
Loss function curve of three models in Tripleknock. **a**, MSE curve of AE1. The x-axis represents mini-batch epochs, accumulated across 10 parts of the training set due to memory constraints. The green dashed line indicated the background random noise. **b**, MSE curve of AE2, the background noise is not needed because the AE2 doesn’t receive the raw data. **c-d**, BCELoss of MLP model trained on 5% and 10% data.

**Supplementary Table 1.** Phylogenetic tree analysis statistics by BV-BRC.

| Parameters | Value |
| --- | --- |
| Requested genomes | 7 |
| Genomes with data | 7 |
| Max allowed deletions | 0 |
| Max allowed duplications | 0 |
| Single-copy genes requested | 100 |
| Single-copy genes found | 100 |
| Num protein alignments | 100 |
| Alignment program | mafft |
| Protein alignment time | 102.9 seconds |
| Num aligned amino acids | 41,482 |
| Num CDS alignments | 100 |
| Num aligned nucleotides | 124,446 |
| Best protein model found by RAxML | JTTDCMUT |
| Branch support method | RAxML Fast Bootstrapping |
| RAxML likelihood | -503079.0903 |
| RAxML version | 8.2.12 |
| RAxML time | 103.3 seconds |
| Total time | 249.7 seconds |

